# Localized axolemma deformations suggest mechanoporation as axonal injury trigger

**DOI:** 10.1101/816108

**Authors:** Annaclaudia Montanino, Marzieh Saeedimasine, Alessandra Villa, Svein Kleiven

**Affiliations:** Division of Neuronic Engineering, Royal Institute of Technology (KTH), Huddinge, Sweden; Department of Biosciences and Nutrition, Karolinska Institutet (KI), Huddinge, Sweden

**Keywords:** Mechanoporation, Axolemma, Axonal injury, Membrane permeability, Traumatic brain injury, Finite Element

## Abstract

Traumatic brain injuries are a leading cause of morbidity and mortality worldwide. With almost 50% of traumatic brain injuries being related to axonal damage, understanding the nature of cellular level impairment is crucial. Experimental observations have so far led to the formulation of conflicting theories regarding the cellular primary injury mechanism. Disruption of the axolemma, or alternatively cytoskeletal damage has been suggested mainly as injury trigger. However, mechanoporation thresholds of generic membranes seem not to overlap with the axonal injury deformation range and microtubules appear too stiff and too weakly connected to undergo mechanical breaking. Here, we aim to shed a light on the mechanism of primary axonal injury, bridging finite element and molecular dynamics simulations. Despite the necessary level of approximation, our models can accurately describe the mechanical behavior of the unmyelinated axon and its membrane. More importantly, they give access to quantities that would be inaccessible with an experimental approach. We show that in a typical injury scenario, the axonal cortex sustains deformations large enough to entail pore formation in the adjoining lipid bilayer. The observed axonal deformation of 10-12% agree well with the thresholds proposed in the literature for axonal injury and, above all, allow us to provide quantitative evidences that do not exclude pore formation in the membrane as a result of trauma. Our findings bring to an increased knowledge of axonal injury mechanism that will have positive implications for the prevention and treatment of brain injuries.

## 1 Introduction

Traumatic brain injury (TBI) is defined as “an alteration in brain function, or other evidence of brain pathology, caused by an external force”.^1^ In 2013 approximately 2.8 million TBI-related emergency department visits, hospitalization, and deaths occurred in the United States.^2^ In a recent study reporting the epidemiology of TBI in Europe, Peeters and coworkers analyzed data from 28 studies on 16 European countries and reported an average mortality rate of ≈ 11 per 100,000 population over an incidence rate of 262 per 100,000 population per year.^3^ Diffuse axonal injury (DAI), a multifocal damage to white matter axons, is the most common consequence of TBIs of all severities including mild TBIs or concussions.^4^ Invisible to conventional brain imaging, DAI can only be histologically diagnosed and its hallmark is the presence of axonal swellings or retraction balls observable under microscopic examination.^5^

Considerable research effort has been put into the understanding of the primary effects of trauma onto the neurons. To define an axonal injury trigger, nervous tissues and neuronal cultures have been subjected to dynamic loads leading to several hypotheses regarding the cell injury mechanism. Mechanoporation (i.e. the generation of membrane pores due to mechanical deformation) of the axolemma has been historically put forward as primary axonal injury mechanism.^6–11^ Although such a mechanism was initially proposed to affect the somatic plasmalemma, it was more recently associated with both the somata and neural processes.^12,13^ Assessing the occurrence of such a mechanism is particularly important in cases of mild/moderate TBIs, where cells’ abnormalities – but not immediate death-are coupled with functional alterations.^14^ Disruption of the axolemma has been inferred observing changes in membrane permeability due to mechanical stretch supposedly leading to increased intra axonal calcium concentration, however this has been questioned by some studies.^15–17^

Experimental studies using oriented cell cultures have instead put forward microtubule disruption as cell injury trigger having observed cytoskeletal damage and tau protein accumulation at the site of axonal swelling.^18,19^ These studies have further motivated the development of models (both analytical and computational) having the microtubule bundle as a focus.^20–25^ In our previous work, however, we have shown with a finite element model of the entire axon that, especially when including detachment of tau protein elements, deformations in the microtubules did not pass the suggested microtubule failure thresholds.^26^ On the contrary, we found that, as a result of axonal stretching and of microtubules distancing, very high strain localization formed on the axonal membrane. However, we were unable to say whether these could lead to axolemmal damage or not.

Up to date, studies assessing poration thresholds – both experimental and computational ones - have not been axon-specific and therefore might not reflect axolemmal behavior. Previous experimental studies have in fact focused on areal deformations of red blood cells (RBCs).^27^ Molecular simulations studies have investigated mechanoporation by deforming two-to-four component lipid bilayer biaxially.^28–30^ Given the slenderness of the axons, however, during stretch-injury one can expect the axolemma to deform mainly along the axial direction. Not only loading mode specificity, but also material specificity might also play a key role in the understanding of axonal injury. Among others, lipid composition is known to affect membranes’ biomechanical properties.^31^ For example, experimental studies have established areal failure thresholds (ε_A_ <10%) based on RBC’s or giant lipid vesicles’ membranes.^32,33^ It has however been shown that these thresholds do not apply to the axons’ membrane: when undergoing osmotic shocks - which induce biaxial deformation of the axolemma-in fact the axolemma could sustain areal strains ε_A_ > 20%) without rupturing.^34^

Another peculiarity of the axonal membrane is that both in unmyelinated and myelinated axons, protein channels span the lipid bilayer, making possible the propagation of the action potential. In particular, voltage-gated sodium channels are the main responsible for the continuous and saltatory conduction in unmyelinated and myelinated axons respectively. The density of sodium channels in the axonal membrane has been reported to be in the range 5-3000 channels/μm^2^. Lower densities of sodium channels are found in unmyelinated axons, while higher densities are observed in nodal portions of myelinated axons. ^35,36^ These channels have also been proposed to influence the injury response.^37^ Rigid inclusions embedded in a lipid bilayer model have been shown to facilitate membrane disruptions while stiffening the membrane-inclusion system.^38^ In this study, unmyelinated axons are considered. These axons are, in fact, not only as numerous as myelinated ones in the human brain,^19^ but have also been shown to be more vulnerable to mechanical strain than their myelinated counterpart.^39,40^ These motivations have so far justified the usage of unmyelinated axons as experimental injury models. Modelling an unmyelinated axon allows us to gain a better insight of the basic axon injury mechanism, irrespective of more complex and ill-defined boundary conditions -such as those that could be induced especially at the paranodal junction by the attachment of myelin or other surrounding tissue.

It is apparent that, in order to describe the axonal injury mechanism, both axon-specific boundary conditions and material properties should be taken into account. Therefore, to provide mechanical insights into the initiation of axonal damage, we combined a finite element (FE) model of a generic distal portion of an unmyelinated axon and a molecular-based lipid bilayer model with fixed number of particles. In particular, the axonal model was utilized to simulate typical stretch injury scenarios (Figure 1a, 1d). As a result of cellular-level deformation, local deformations happening at the cortex level (Figure 1b, 1e) could be extracted. These local deformations were then used as input for molecular simulations of the axonal lipid bilayer (Figure 1c, 1f). In this way, membrane permeability or poration could be quantified in dependence of applied axonal strain and strain rates.

**Figure 1:**
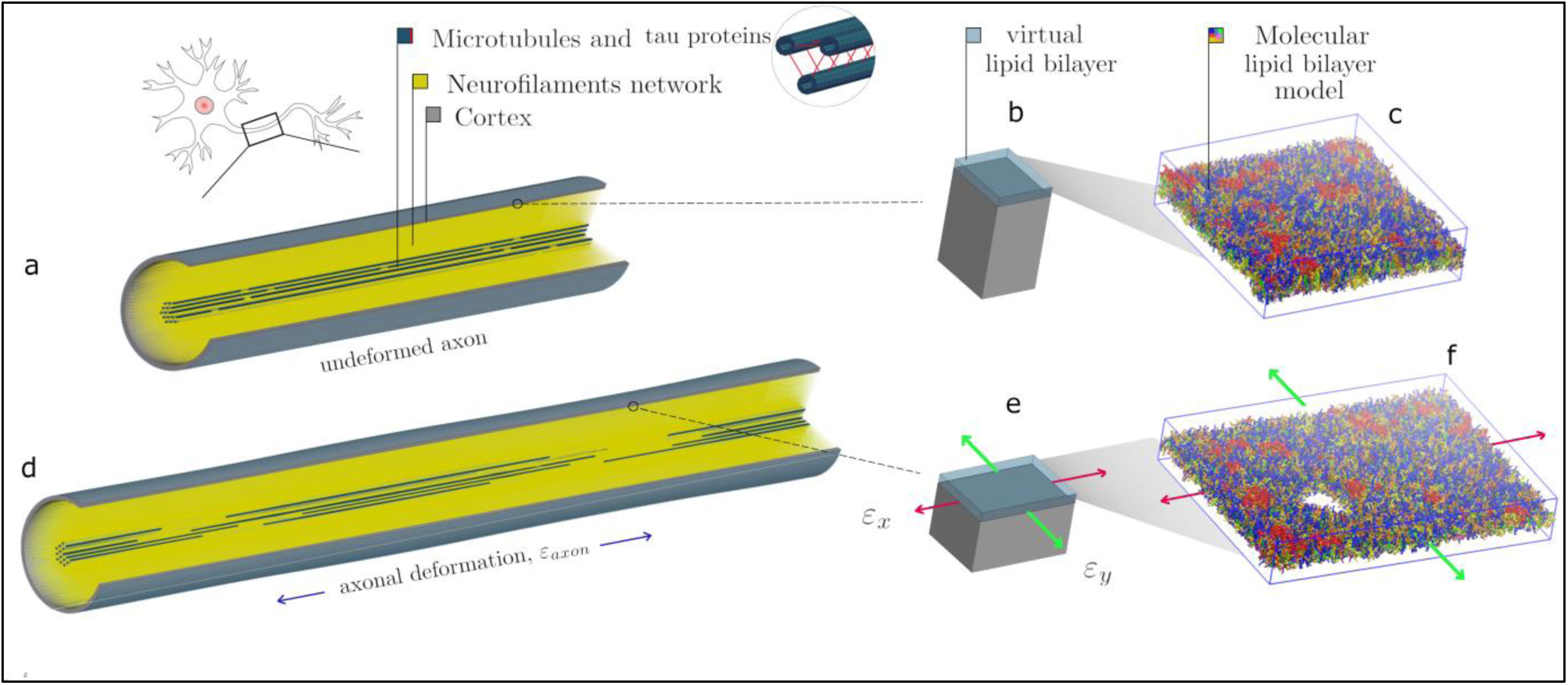
Schematic representation of the combined modelling approach. **(a)** Finite element axonal model coated with a slightly transparent layer symbolizing the lipid bilayer (virtual). A quarter of the model was removed to reveal the arrangement of the inner structures. A microtubules bundle (solid blue hollow cylinders cross-linked by beam-like tau proteins) is located centrally, while the rest of the axonal space is filled with a dense beam-meshed neurofilaments network (yellow). The cytoskeleton is wrapped in a thick shell layer representing the axonal cortex (grey) **(b)** Single undeformed cortex element coated with a virtual lipid bilayer linked to the corresponding undeformed molecular coarse grained model of the lipid bilayer **(c)**. The lipid content resembles the plasma membrane: (PC lipids are in blue, PE in yellow, CHOL in green, SM in orange, PS in purple, GM in red, anionic lipids in pink, and the rest of lipids in gray, water is not visualized for clarity). **(d)** Deformed axon finite element model (axonal deformation, ε_axon_) **(e)** Deformed cortex element (localized deformation: ε_x_=0.34, ε_y_=0). **(f)** Visualization of the induced pore upon lipid bilayer deformation.

## 2 Methods

### 2.1 Finite element (FE) framework

FE head models are commonly used by the scientific community, despite their level of idealization, for the explanation they can provide of traumatic phenomena. Through FE simulations it is in fact possible to simulate traumas and extract quantities that would otherwise be inaccessible via an experimental setup. Similarly, the FE method has been used to capture the mechanical behavior of cells and subcellular components.^41–47^ More recently, FE models of the axon have been proposed to shed a light on the mechanism behind axonal injury.^25,26,48–50^

In the present study a previously published and validated axon FE model was used (Figure 1a), which is a 8 μm long representative volume (RV) of an axon consisting of three main compartments: a microtubule (MT) bundle, the neurofilament (NF) network and, finally, the axolemma-cortex complex wrapping the entire structure.^26^ The axon model diameter (1.15 µm) was chosen to fall in the range of those from experiments to which it was originally validated against. ^51–53^ The choice of its length was instead dictated by considerations related to the average MT length. Given the average continuous length of MTs (4.02 µm),^54^ to contain a unit length of the microtubule bundle the length of the axon RV was set to 8 µm. Given its cross-section and the proposed MTs densities, 19 rows of MTs were included in the axon RV, each containing two randomly placed discontinuities.^55–57^

In the MT bundle, solid elastic MTs are cross-linked by viscoelastic tau proteins, which are modeled as discrete beams. These are assigned a failure threshold so that they stop carrying the load as soon as they reach double their length.^20^ The NF network is modeled as a dense mesh of viscoelastic beams filling up the axonal space and anchoring the MTs to the plasma membrane.^58^ A detailed description of our literature-supported modeling choices can be found in the supplementary material and in our previous publication.^26^

In a study focused on the assessment of RBC membrane mechanical behavior, Evans and coworkers defined the lipid bilayer as "along for the ride" when the cell is deformed, meaning that the cortex (or “matrix”) is the one resisting the applied deformation and influencing the structural response.^59^ Assuming this to be true also for the axon, in the current study, the axonal wrapping layer was assigned properties in line with those of the sole axonal cortex. The axonal cortex has a unique periodic organization made of rigid actin rings connected by flexible cylinders of spectrin.^60^ In our model the cortex is represented by a layer of fully integrated shell elements with a thickness of 50 nm.^61^ These elements were assigned a shear modulus G = 0.0016 MPa and a viscosity η =10^5^ Pas.^62,63^ Mechanical properties of all the model components can be found in Table 1. In a recent study, fluid friction was observed at the membrane-cortex interface with friction forces being proportional to the relative speed of the two layers.^61^ Considering the axonal loading mode chosen for this investigation (i.e. pure stretching), however, the relative movement between the two layers can be assumed to be minimal. Hence, we assumed the lipid bilayer as fully tied to the cortex: any deformation happening in the cortex plane would be directly transferred to the lipid bilayer. Therefore, in the current study an explicit FE representation of the membrane was avoided (hence the "virtual" representation in Figure 1) and deputed instead to a molecular dynamics description. In practice, cortical deformations resulting from axonal injury simulations were extracted and fed into coarse grained (CG) membrane or membrane-protein systems.

**Table 1:** Axon model material parameters. Young+s modulus (E), Poisson’s ratio (ν), shear modulus (G), viscosity (η), spring stiffness (K), 1D viscosity coefficient (μ) (for references, see supplementary material)

| Axonal component | Element type | Material |
| --- | --- | --- |
| Microtubules | Solid | Linear elastic<br>$E=830$ MPa<br>$\nu=0.37$ |
| Cortex | Shell | $G=0.0016$ MPa<br>$\eta=1e5$ Pas |
| Tau proteins | Discrete beam | Linear viscoelastic<br>$K=5e-5$ N/m<br>$\mu=2.205e-4$ Ns/m |
| NFs elements | Discrete beam | Linear viscoelastic<br>$K=7.5e-5$ N/m<br>$\mu=1e-7$ Ns/m |
| Mts-NFs links | Discrete beam | Linear viscoelastic<br>$K=1e-6$ N/m<br>$\mu=1e-7$ Ns/m |

### 2.2 Axonal injury simulations

To study axonal injury at the single-cell level it was deemed appropriate to consider the axonal behavior under uniaxial deformation. It is generally accepted that strain is the main mechanism behind axonal injury.^64^ Although at the tissue (i.e. brain) level strain presents itself as a 3D tensor, to enforce experimental control, it is common to apply only one deformation at a time, this being of compressive, tensile or shearing nature. It has previously been shown with neuronal cultures that different loading modes might actually lead to different mechanisms of injury.^65^ Specifically, in that study, Geddes-Klein and coworkers found that stretching a primary cortical neuron culture uniaxially or biaxially (simultaneously in two perpendicular directions) yielded different mechanisms behind the influx of calcium. However, it should be considered that when deforming a network of cells - which are not embedded in a matrix-, the network’s almost 1D structures, the single axons, will first re-orientate in the direction of the load and then mostly stretch.

In an effort to investigate a general axonal behavior, rather than a particular one, 10 different FE axon models were produced by altering the MT bundle geometry of the baseline model. More specifically, every model was generated by randomly moving the MTs discontinuities locations, while keeping the average MTs length of the original model. Symmetry conditions were enforced on a quarter of a model and simulations were run with the latter condition.

Each axon FE model was subjected to a displacement-controlled uniaxial tensile deformation (Figure 1d). Namely, each model was stretched up to a global strain ε_*axon*_ = 30%. This deformation range was chosen to cover all the axonal injury thresholds proposed so far in the literature at either cellular or tissue level.^66–70^ The deformations were applied at strain rates 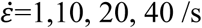, which are characteristic of axonal injury experiments.^7,9,18^ It is important to note that our axon FE model represents an unmyelinated axon and hence can be compared against *in vitro* injury model, which typically do not present myelinated axons. All simulations were performed in LSDYNA using an implicit dynamic solver. The implicit scheme was a necessary choice dictated by the element dimension, while the dynamic regime was chosen to be able to capture strain-rate related inertia and material effects. Subsequently, for each simulation of 1,10, 20 and 40 /s strain rates the 1^st^ and 2^nd^ principal strains (*ε*_*x*_,*ε*_*y*_) in the cortex plane were extracted as function of axonal strain.

### 2.3 Setting up the membrane molecular model

The membrane was modeled as a molecular-based lipid bilayer (two leaflets of lipids) and was described by a CG model where groups of atoms (3-4 heavy atoms) are united into beads which interact with each other by means of empirical potentials. The MARTINI2.2 force field together with the non-polar water model was used.^71–73^ The axolemma is characterized by having phosphatidylcholine (PC), phosphatidylethanolamine (PE), cholesterol (CHOL), glycolipid, (and for some studies also sphingomyelin (SM)) as the main components (**Table S1**). To mimic the axolemma’s lipid composition, the mammalian plasma membrane model deposited on MARTINI webpage (http://www.cgmartini.nl/) was used.^74^ In the membrane model, 63 different types of lipids were distributed asymmetrically between two leaflets (Figure 1c). The outer leaflet has a higher level of saturation of the tails and contains PC (36%), PE (6%), CHOL (31%), SM (19%), glycolipid (GM) (6%), and other lipids (2%). The inner leaflet, which has a higher level of polyunsaturation, contains PC (17%), PE (25%), CHOL (29%), SM (9%), phosphatidylserine (PS) (11%), anionic lipids phosphatidylinositol (PIP) (2%), and other lipids (7%). Note that the reported values are in mol %. Full details of the membrane model can be found in the work by Ingolfsson and coworkers.^74^ The bilayer (containing a total of 6700 lipids) was placed in a cubic box (42*42*12 nm) and solvated by about 100,000 CG water beads and NaCl was added to mimic the ionic strength at physiological condition (150 mM NaCl).

### 2.4 Setting up of membrane-protein molecular model

Sodium channel protein type subunit alpha (Na_v_1.1) was used to represent the family of voltage-gated ion channels embedded in the axonal membrane.^75^ The three-dimensional (3D) structure for the homo sapiens Na_v_1.1 has not been resolved experimentally, however, the encoding gene is known (SCN1A gene).^76^ Thus we built the 3D structure using homology modeling and Phyre2 server was used.^77^ Among the available sodium channels, the best sequence alignment (100% confidence and 48% sequence identity) was found with putative sodium channel from American cockroach, Na_v_PaS (PDB ID: 5X0M).^78^ The cryo-electron microscopy structure of Na_v_PaS has been used as template. The missing N-terminal and C-terminal domains, 1-2 and 3-4 linkers were modeled using the software Modeller.^79^ The so obtained 3D structure for Na_v_1.1 is composed of a single polypeptide chain that folds into four homologous repeats (Figure 2), each one containing six transmembrane segments. The protein was oriented in the lipid bilayer using Positioning of Proteins in Membrane (PPM) method.^80^ The structure was first minimized in the PC bilayer at the atomistic level (using CHARMM36 force field^81^) using 5000 steps of steepest descent minimization and then equilibrated for 200 ns.

**Figure 2:**
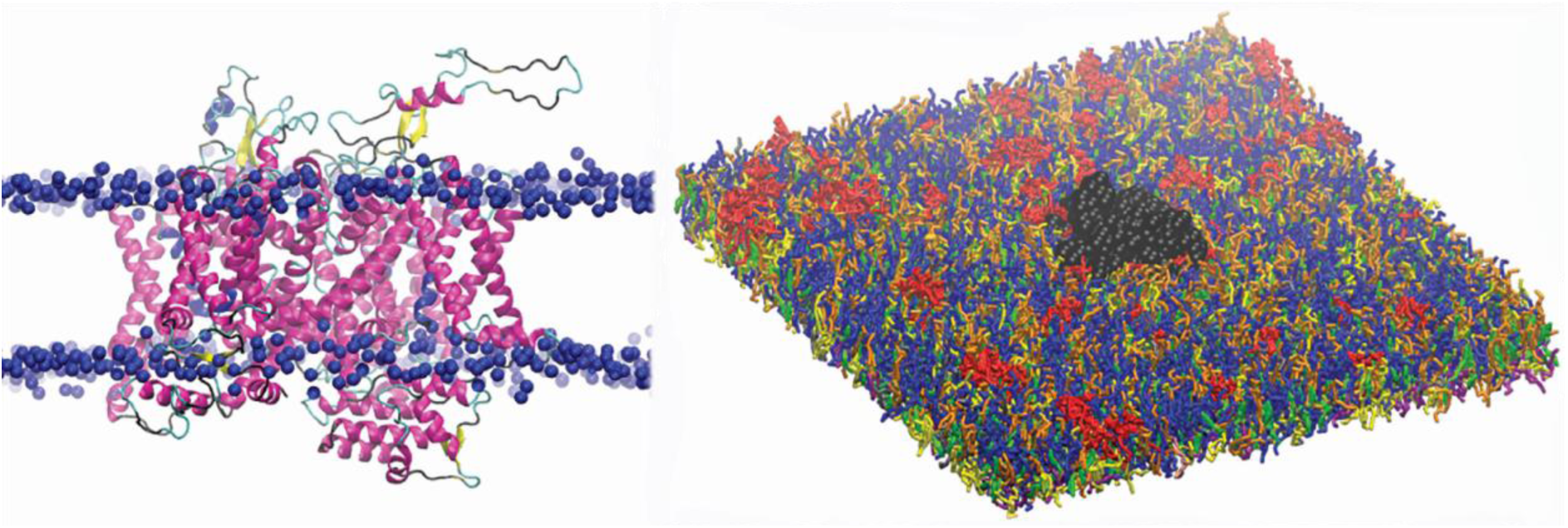
(a) Three dimensional structure of Na_v_1.1 protein embedded in the lipid bilayer. Protein is shown by secondary structure elements (α-helix in magenta, β-sheet in yellow, turn in cyan, and coil in black). Phosphate atoms of lipid head group are shown in blue. (b) CG model of Na_v_1.1 protein in the plasma membrane model. Protein is shown in black VDW representation. For lipid color code see Figure 1.

The obtained atomistic structure was used to build the protein CG model. Martinize protocol together with ElNeDyn were used to convert the atomistic Na_v_1.1 structure into a coarse-grained model.^82,83^ One protein was embedded in the 42×42 nm^2^ CG membrane (Figure 2), that corresponds to a density of 567 ion channels/μm^2^.

### 2.5 Molecular Dynamics (MD) simulations

MD simulations are widely used to provide a three-dimensional description of lipid bilayers and proteins at the molecular level and their evolution in time, from which time-dependent and independent properties can be evaluated.^84^ All MD simulations were performed using the GROMACS simulation package, version 2016.^85^ Simulations have been performed in NPT ensemble (constant number of particles, constant pressure, and constant temperature) and NP_z_AT ensemble (constant number of particles, constant pressure along z, P_z_, constant bilayer area, and constant temperature). The temperature of the systems was kept at 310 K using velocity rescale thermostat with a time constant of 1ps.^86^ Semi-isotropic Parrinello-Rahman barostat was applied to couple the pressure to 1 bar with a time constant of 12 ps (compressibility of 3e-4 bar^−1^).^87^ Periodic boundary conditions were applied. A time step of 20 fs was used. The Verlet cutoff scheme was used. Non-bonded interactions were calculated using a cut-off of 1.1 nm.^88^ The reaction field potential was used to treat long range electrostatic interactions using a switching distance of 1.1 nm.^89^

NPT simulations were performed to equilibrate the membrane structure in absence of deformation. To generate the starting structures for deformed lipid bilayers, we deformed the bilayer using unsteady stretching algorithm with a stretching speed c=5 m/s. The so-obtained deformed system was then simulated at constant bilayer area for 2 µs to generate an ensemble of configurations that describes the membrane at the selected strain. The simulations were extended up to 5 µs for strains larger than 30 %. During the simulations an elastic bond force constant of 500 kJ.mol^−1^.nm^−2^ was applied to the protein atom pairs within a 0.9 nm cut-off (ElNeDyn).^83^ That means that the protein is simulated as a semi-rigid body. The analysis was performed on the last 1μs of production runs. The errors were calculated by block averaging over 5 blocks. All the figures were rendered using Visual Molecular Dynamics software.^90^ As criteria to detect pore formation in the lipid bilayer, we use the jump in the surface tension. After the detection of the pore, simulations were extended of 5 to 9 µs to check whether spondaneous sealing occurred in the simulation time scale (microseconds).

### 2.6 Permeability coefficient

The transport of substances across a lipid membrane is a biological process of vital importance. Small molecules, such as water molecules or drugs, can be transported into the membrane in a passive or active way. A passive transport proceeds via an entropy-driven, nonspecific diffusion process of the molecule across the membrane. Permeability or leakage of a small molecule should give a measure of the structural stability of the membrane since it should reflect lipid packing at the membrane core. Membrane permeation can be described by a so called "solubility-diffusion mechanism" (see review by Shinoda^91^ for details) and quantified by the membrane permeability coefficient, *P*_*m*_. The *P*_*m*_ *of a* water molecule is proportional to its oil/water partition coefficient (*K*_oil/water_).^92,93^

First, we calculated the *membrane/water* partition coefficient for water at equilibrium and at different strains. We do that by calculating the average number of water molecules (1 CG water bead = 4 water molecules) in the bilayers within a distance of 0.52 nm from the lipid tails on 1 µs molecular simulations. The partition coefficient for water is 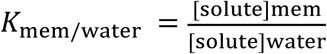, where [solute] denotes the concentration of water molecules in the lipid bilayers and water solution, respectively. [water]_mem_ is obtained by dividing the average number of water molecules by the lipid bilayers volume, while for the water concentration in water the experimental value (55.5 mol/L) was used. The membrane volume was corrected to account for the presence of the protein. Knowing that log*P*_m_ is proportional to log*K*_mem/water_, we then estimated how membrane strain influences its permeability.

## 3 Results

In this study, axonal injury was simulated deforming a FE model of the axons of a quantity ε_*axon*_. Cortex principal strains (ε_*x*_,ε_y_) were then extracted and applied in cascade to a molecular model of the plasma membrane and the structural outcome on the latter was observed. As evidenced by Figure 3 - which shows the distribution of 1st Principal Green Lagrange strains along the cortex shell-layer for one of the 10 tested models for 5% < ε_*axon*_ <15%-when the axon is stretched (0< ε_*axon*_ <30 %), due to its composite nature, the deformation pattern along the axonal cortex is not homogeneous. Areas of strain concentration on the cortex evidently form throughout axonal deformation. The magnitude of this strain concentration is dependent both on the applied strain (different columns in Figure 3) and strain rate (different rows in Figure 3). It can be observed that, for example, an applied axonal strain as low as 5% corresponds to a local maximum strain which is twice as high. The more ε_*axon*_ increases, the more this localization of strains in the cortex is pronounced. Moreover, on each column it is possible to observe, for the same axonal strain, the influence of strain rate (1, 10 and 40 /s) on the local deformation level. Here it is possible to notice that higher strain rates do not lead to higher local maximum strains for ε_*axon*_<10%. However, at higher axonal strains the intuitive order is re-established: higher strain rates lead ultimately to higher cortex-level deformations (**Figure S2**).

**Figure 3:**
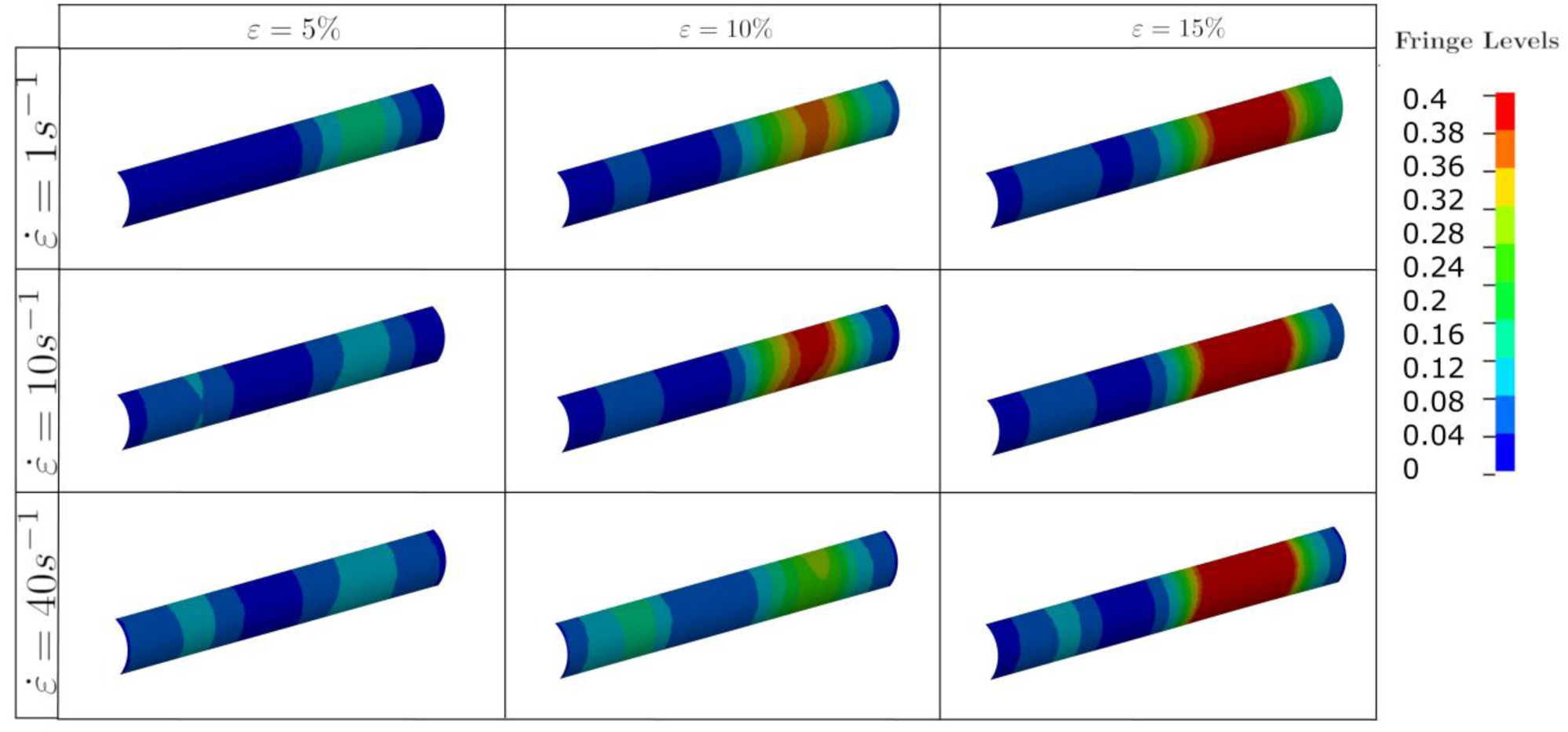
Fringe plots showing the 1st Principal Green-Lagrange strain along the axonal cortex as a result of 5, 10, and 15 % axonal strains in the first, second and third column respectively. In the first, second and third row results for strain rates of 1, 10 and 40 /s are reported. The range was set between 0 and 40% for visualization purposes.

To better understand this behavior, the average results of 10 different models are reported in Figure 4 together with the standard deviations for each tested strain rate. In this plot the cortex maximum 1^st^ Principal Green-Lagrange strains (ε_x_) are reported as a function of the applied axonal strains. What can be noted is, first, the increased standard deviation with higher rates. In addition, one can notice that for strain rates of 1 /s and 10 /s the curves follow the same path until ε_axon_ ≈ 7% and then diverge. Interestingly, as previously noted, the response for a strain rate of 40 /s is on average lower than that of 1 /s and 10 /s until ε_axon_ ≈ 12% and ε_axon_ ≈ 20% respectively. 2^nd^ Principal Green-Lagrange strains (ε_y_) in the cortex plane were consistently found to be at least 6 orders of magnitude inferior to ε_x_, therefore they were set to zero when input into MD simulations.

**Figure 4:**
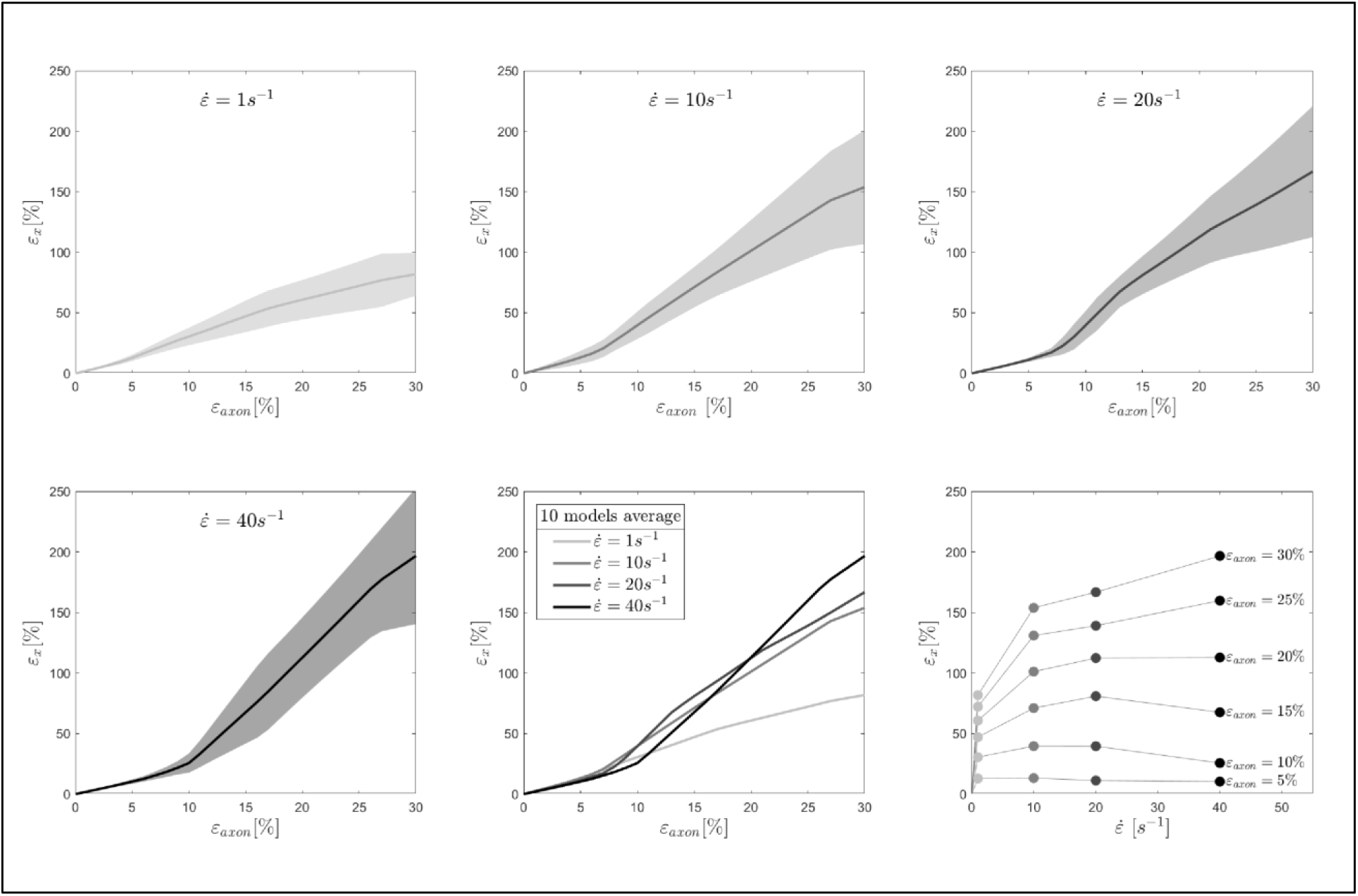
Maximum 1st Principal Green-Lagrange strains in the axonal corex as a function of axonal strain. Lines represent the average over 10 models and the shaded areas represent the standard deviations. Results are shown for strain rates 1,10, 20, and 40 /s. In the second row (center), the means for the different strain rates are shown in the same plot to ease the visual comparison. Finally, in the second row (right), iso-strain curves are reported, which show the maximum 1st Principal Green-Lagrange strains in the cortex as a function of the applied strain rate.

To understand the effect of axonal deformation on the membrane integrity, we used MD simulations and a molecular based-lipid bilayer to describe the axonal membrane. First, to verify that the membrane models resemble the mechanical feature of natural membrane, the area compressibility modulus was estimated for small areal strain values (less than 0.05). A value of 303±6 mN/m and 352±3 mN/m, respectively in absence and presence of Na_v_1.1 protein, was obtained (**Figure S3**) – which is in agreement with the area compressibility modulus measured for red blood cell membranes (375±60 mN/m) at 37°C by Waugh and Evan.^94^ The agreement allows us to move forward and use the model to investigate the effect of local deformation on the bilayer structure. A selected number of cortex strains ε_x_ (Figure 4) were input into the membrane and membrane-protein system, while the membrane model was kept undeformed in the y-direction (ε_y_=0).

We monitored the bilayer structural features (i.e. pore formation, interdigitation, and water permeability) at different local strains. Figure 5 depicts how the occurrence of pore formation is detected. Namely, this event corresponds to a jump in the surface tension. Figure 6 summarizes which molecular-level event is to be expected at each axonal strain ε_axon_ (and related maximum cortex strain, ε_x_) for a plasma bilayer model with and without embedded protein. The lipid bilayer can withstand the applied strain value of ε_x_ = 34%, corresponding to ε_axon_ = 10-12 %. At higher strains pore formation is observed. When Na_v_1.1 is embedded in the membrane, poration occurs at a larger cortex strain (ε_x_ = 47 %), corresponding to ε_axon_ = 12-15 %.

**Figure 5:**
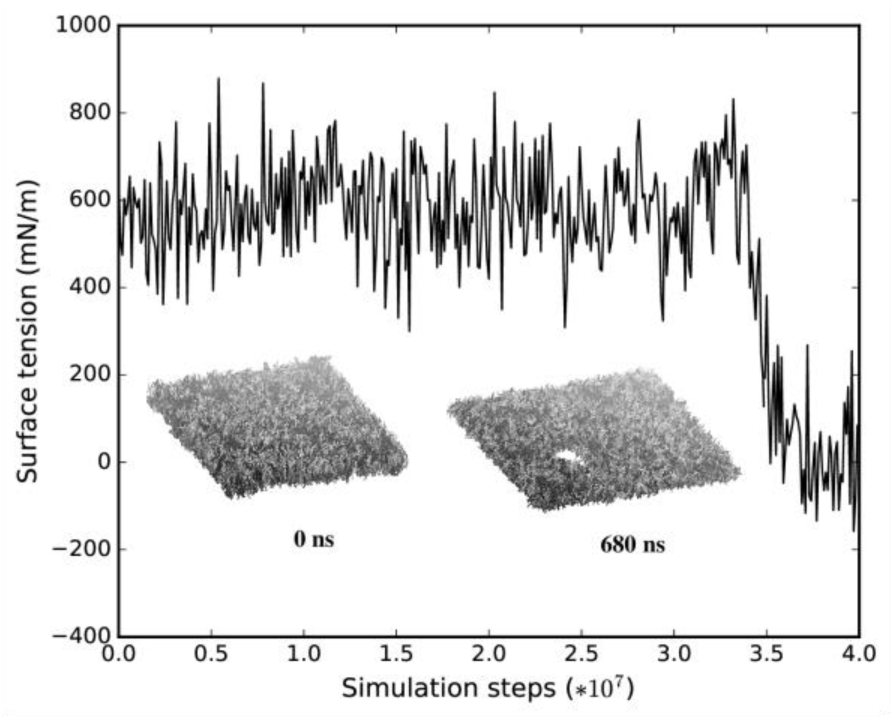
Surface tension and corresponding lipid bilayer structure at ε_x_=0.34 as a function of simulation steps.

**Figure 6:**
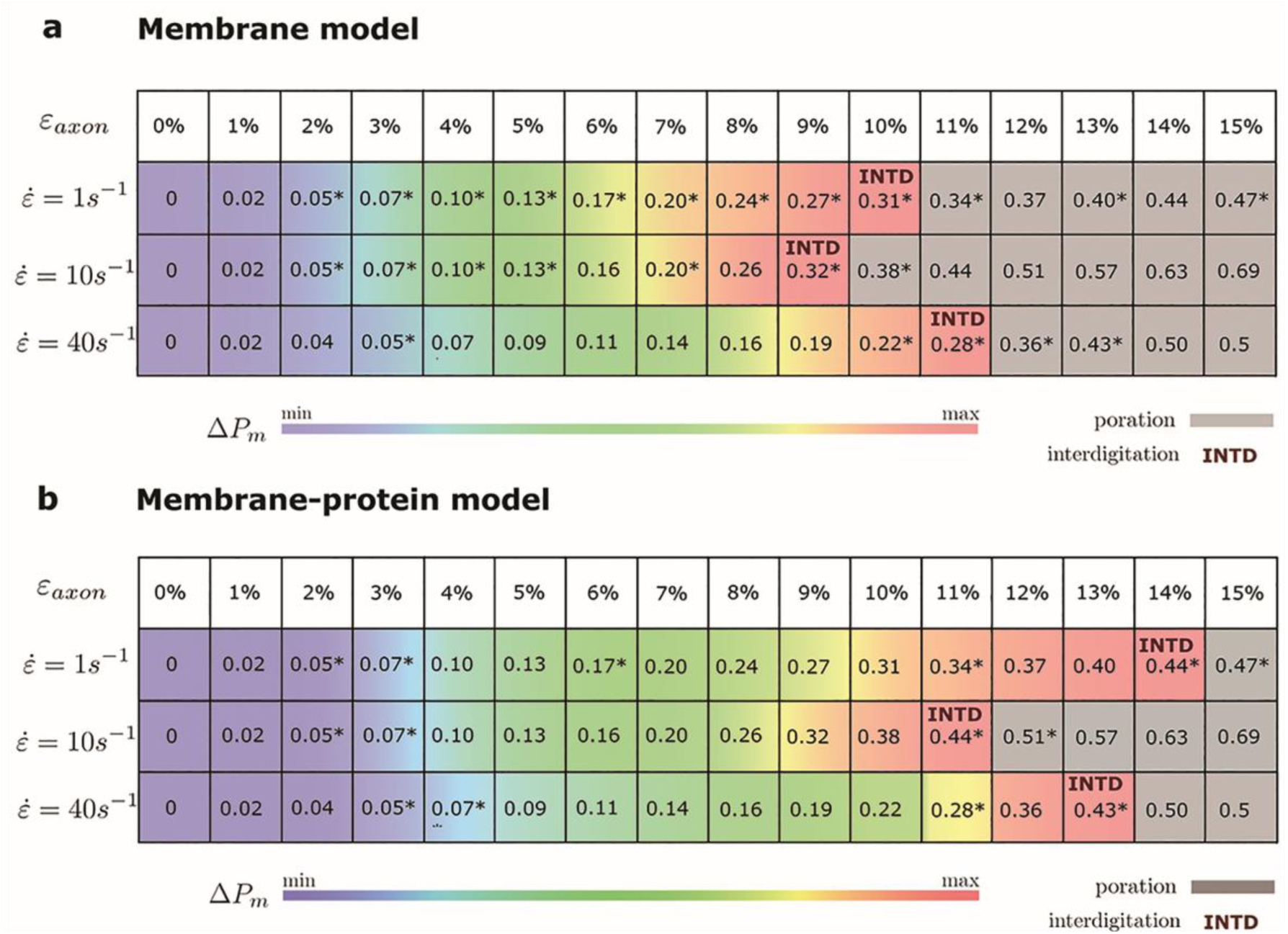
Axonal strains and corresponding maximum local strains (mean values over a family of 10 different FE axonal models) together with strain rates. * indicates the local deformations for which molecular dynamics simulations have been performed. Top panel refers to the membrane model and bottom panel to the membrane-protein model. Δ*P*_*m*_ indicates how water permeability increases with respect to the equilibrium value (at local strain=0). max corresponds to 40 % in the membrane model and 51% in the membrane-protein system.

Figure 7 shows an example of the observed bilayer disruption in absence and presence of Na_v_1.1 protein (**MovieS1**, **Movie S2**). We observed that the membrane rupture occurs at slightly larger local strain in presence of the protein. The presence of the Na_v_1.1 protein therefore not only renders the membrane more resistant to deformation but also to rupture. In addition, it was observed that before poration, the bilayers start to form interdigitated states (**Figure S4**). Interdigitation occurs at ε_x_ > 27 % in absence of protein and ε_x_ > 34 % in presence of protein, corresponding to ε_axon_ =9%-11% and ε_axon_ =10-12%, respectively (Figure 6). Interestingly, poration both in presence and absence of protein takes place in bilayer regions lacking ganglioside lipids (Figure 7). Ganglioside-type lipids are mainly found in the outer leaflet of the membrane and are known to make the lipid bilayer more resistant to deformation due to the interactions between sugar headgroups.^30^

**Figure 7:**
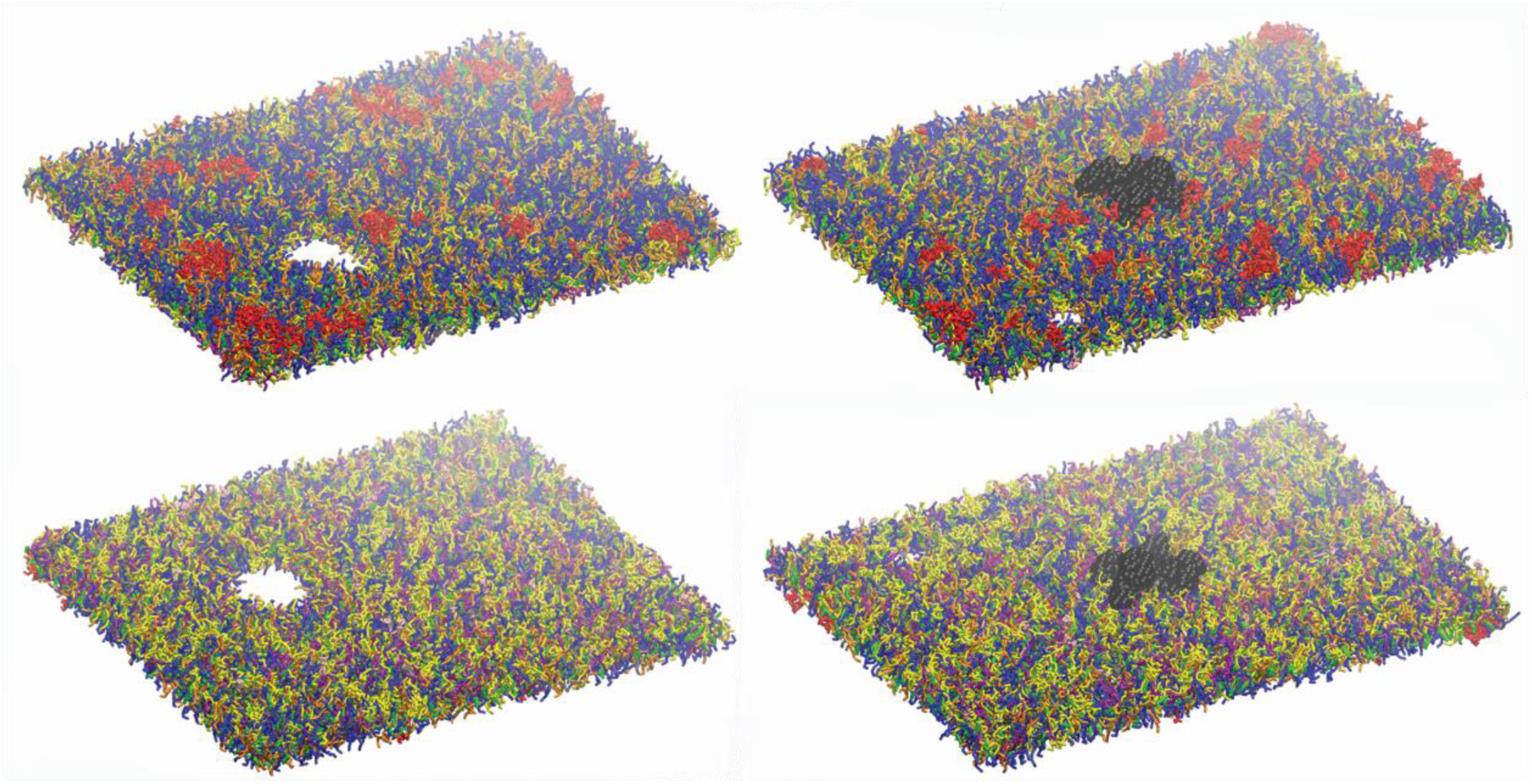
Lipid bilayers disruption. Left panels: snapshot at 680ns at ε_x_=0.34 in absence of Na_v_1.1. Right panels: snapshot at 27 ns for ε_x_=0.47 in presence of Na_v_1.1. (in black). Top panels show the upper leaflets, low panels show the lower leaflets. For lipids color code see Figure 1.

To quantify that the molecular models indeed reproduce membrane partitioning, we calculated the membrane/water partition coefficient for water at equilibrium (log*K*_mem/water_ = - 1.73). The presence of the protein has almost no effect on the partition coefficient (log*K*_mem/water_ = −1.75). At the increase of ε_axon_, water partitioning between aqueous solution and lipid bilayers increases proportionally to ε_x_ (**Figure S5**). Given the aforementioned proportionality between the partition coefficient *K*_mem/water_ and the permeability *P*_m_, the latter can be quantified as a function of strain. In particular, starting from equilibrium state, water permeability increases up to 40 % (51 % in presence of protein) before pore formation. Figure 6 coloring describes the change in water permeability for the the lipid bilayers with and without protein channel at the increase of ε_axon_ for different strain rates.

## 4 Discussion

The biomechanics of TBI has been the focus of considerable research effort for many years now. Nevertheless, the underlying injury mechanism leading to cellular impairment is yet to be explained. The use of *in-silico* models can provide better understanding of the mechanical cues determining cells’ condition immediately after a mechanical insult. In this study, for the first time, we were able to provide direct mechanical evidence of the possible onset of mechanoporation as a result of the mechanical insult. More specifically, bridging finite element and molecular dynamics simulations, we could determine at which level of axonal strain the lipid bilayer sustains local deformations high enough to induce pore formation.

Several studies so far have stressed the need of taking into account axonal direction and kinematics when studying injury at the tissue level both in experimental set-ups and computational models. ^95–97^ It has indeed been assessed from tissue or cell cultures that deformations applied to a tissue do not necessarily coincide with those sustained by the embedded axons. In fact, in both tissue models -such as the optic nerve or organotypic slices- and cell cultures non-affine deformations might arise. Similarly, at the cellular scale, which is the focus of the current study, the applied axonal strains ε_axon_ do not directly transfer to the axonal membrane, due to the axonal composite nature. Despite being often assumed as a homogeneous viscoelastic entity, the axon presents dramatic heterogeneities in deformation along its length when stretched.^98^ Evidence of this is also the localized axonal length increase (up to 65%) that was observed as a result of axonal stretch.^19^ The morphological response to stretch injury – such as the appearing of undulations and periodical swellings– clearly hints at an heterogeneity in the axonal response to tensile injury loads. This is confirmed by the localization of membrane strains that we observed with our model (Figure 3). This localization is directly related to the point of weakest connectivity of the microtubule bundle.^24,99^

Especially when considering that our axonal finite element model is just an 8 μm-long representative volume of the axon, one should imagine such a strain localization taking place periodically along a periodical repetition of our model. The periodical swelling, which is characteristic of axonal injury, could therefore be explained by the heterogeneous deformations that are revealed by our model.

Recently, an experimental work has tried to quantify axonal membrane strains induced by the deformation applied to a cell culture as a whole.^100^ Differences between applied strains and membrane strains were reported. However, it must be noted that these discrepancies may mostly be due to the non-alignment between applied load and axons’ directions within the culture. Moreover, membrane strains were calculated from the level of separation of fiducial membrane markers whose initial distance is not reported. Hence, a direct comparison between our results and these data cannot be made.

In our results maximum cortex strains clearly appear as a nonlinear function of axonal strain (Figure 3). Less intuitive is however the relation with strain rate. Due to the viscoelastic (and hence rate-dependent) properties of tau proteins and of the cortex itself, at axonal strains lower than 12%, the cortex deforms less at strain rate 40 /s rather than at 1 /s or 10 /s. Similar observations were reported in previous studies based on numerical and analytical models of the sole microtubule bundle.^24,25,99^ In those studies, simulations of the microtubule bundle undergoing stretch revealed a higher resilience of the microtubule bundle at higher stretch rates. Whether this emerging property is reflected by experimental models of axonal injury has, to the best of our knowledge, not been established yet.

To assess the influence of the chosen material properties on maximum cortex strains and on this rate-related phenomenon, a sensitivity study was conducted with one of the ten geometries (**Figure S1** in Supplementary material). The results indicate that, while neurofilament and cortex properties minimally affect cortex strains, tau protein viscosity does, particularly at rate 40 /s. Changes in tau proteins’ viscosity yield similar results at rate 1 and 10 /s. However, at rate 40 /s increasing or decreasing the viscosity of 100%, delays or anticipates, respectively, the crossing of the mechanoporation threshold. A purely elastic behavior for these elements (null viscosity) re-establishes the intuitively expected order: cortex maximum deformation increases with increasing rate. Although the differences by means of maximum cortex strains are minimal, a rate dependent or stress-based failure criterion for the tau protein (or of other filaments) could be more appropriate and might alter this non intuitive relation between poration and strain rate. Alternatively, a surface tension-based poration criterion could be investigated.

Local deformations of the axonal cortex were used as input to MD simulations of the lipid bilayer to establish model-informed axonal injury thresholds. Combining the axonal and membrane level we show that poration occurs in a range of axonal strains going from 10% to 15%. These values are in agreement with previously established axonal injury thresholds.^18,66–70^ Thresholds obtained from experimental tissue injury models cover a slightly higher range of axonal strains.^68,69^ However, this is probably due to the influence of axonal tortuosity and the alternation between an affine/non-affine regime.^95,96^ Namely, when deforming the tissue experimentally, at least initially, part of the tissue deformation goes into straightening of the axons. Only later the axons themselves sustain a deformation.

Several studies have so far tried to assess axonal injury by means of permeability.^6,9,10,12,16,101^ Cell-impermeant macromolecules, such as horseradish peroxidase or Lucifer Yellow, are commonly injected in the extracellular space, prior to the application of a mechanical insult to cell culture or the organ itself. The presence of these molecules in the intra-cellular space post-injury has been proposed as an indicator of pore formation in the axolemma. These changes are studied as a function of the magnitude of the applied strain and strain rate. However, most often results are representative of the entire culture (consisting of non-aligned axons) and not of the single cell. Hence, they cannot be compared with our results that instead relate axonal strains to permeability changes in a localized membrane volume. In a study by Kilinc and coworkers,^10^ however, permeability was observed in individual axons. In particular, membrane permeability in injured axons was found to be twice as high as in control axons. This is in agreement with our results showing that changes in water permeability can be expected to be higher than 40 % (51 % in presence of embedded proteins) at the onset of poration. Nevertheless, our results show that changes in permeability can be expected even below the poration threshold, namely at axonal strains lower than 10-15 %. Interestingly, in a previous study by Yuen and coworkers,^70^ which utilized an aligned cell culture system, pathological axonal alterations were observed at strains starting from ε_axon_=5% – a value that is more than twice as low as established axonal injury thresholds.^18,66–69^ Based on our molecular simulations insight, this might be indeed justified by alterations in axolemma permeability pre-poration.

At this stage, we would like to stress that it is only the combination between these two models (the axonal and molecular-based membrane model) that allowed us to achieve such a good agreement with experimental injury thresholds. In particular, ignoring the composite nature of the axonal model, one would be tempted to assume that the strains sustained by the membrane are equivalent to the tensile strains sustained by the axon. In this scenario, according to our results (Figure 6), poration would occur at ε_x_=ε_axon_ ∈ [0.34,0.51]. These strain values are considerably higher than those so far proposed for axonal injury. Our modeling effort was first aimed at an accurate representation of the axon passive mechanical response to provide axon-specific boundary conditions to simulate molecular-level membrane simulation. Secondly, an accurate representation of the membrane lipid content was deemed necessary to assess features such as membrane permeability and poration given that lipid content is known to have an influence on membrane’s mechanical properties.^28,30,31,33^

Notably, considering the strain concentrations revealed by the axonal model with a different lipid bilayer model would have yielded different results. Pure phospholipidic bilayers, in fact, have been reported not to rupture until 0.68 von Mises strain.^29^

## 5 Limitations

Although our modeling approach brings new insights into the axonal injury mechanism, here we want to address some limitations. Firstly, it must be noted that no active molecular mechanism was considered in our axonal injury simulations. This was deemed a fair approximation given that the aim of this work is to assess injury events that happen in fractions of seconds, whereas biomolecular mechanisms, such as polarization/depolarization, neurofilaments transport etc. live in the *seconds-*time scale. In light of this, our results are to be seen as insights into the possible mechanistic events related to primary injury, rather than damage evolution towards secondary axotomy.

It is also important to stress that our model represents a generic portion within the distal axon of an unmyelinated neuron. Several studies have so far proposed specific axonal sites such as the nodal, paranodal, internodal segments as primary sites of injury.^102–104^ Few studies have also highlighted the susceptibility to injury of the axon initial segment (AIS), the parasomatic region where action potentials are initiated.^105–107^ Due to the differences in the cytoskeleton of the AIS and the distal axon, however, our results cannot be generalized to this segment.

Microtubules failure was discarded as injury trigger in our previous publication.^26^ Nevertheless, primary effects on other filaments might still subsist. For example, neurofilament (NF) compaction was reported to occur as a direct effect of the mechanical insult by Meythaler and coworkers.^108^ However, a larger body of evidence suggests NFs compaction to be associated it with axolemmal permeability changes.^109–111^ Our model includes a representation of the NFs network, whose morphological and material characteristics are reported in the supplementary material. Nevertheless, we are persuaded that the current representation of this network in our FE model, despite being reasonable as a bulk, cannot reveal local phenomena related to injury. The lack of experimental information in fact prevented us from assigning specific properties to filament backbone and side-arms as well as failure properties for their connections. Hence, a priori, we cannot exclude that damage to this component of the model takes place.

At this point, it is to note that our model accounts for the effect of an embedded protein on the membrane structural features, but does not account for possible protein structural change at different strains, since the protein is described as a semi-rigid body. Na_v_1.1 was chosen as a representative example of one of the axolemma ion channels, however our protein-membrane system does not account for the inhomogeneity of proteins distribution along the axolemma. Future studies could aim at assessing the potential effect of protein deformability, compositions or distributions on axolemma stretch susceptibility.

Moreover, the molecular systems used in this study do not account for the recruitment of lipids, which has previously been discussed as a strategy put in place by the neurons to reduce the build-up in membrane tension. ^34,112^ Such a mechanism was however observed in experiments applying a “gentle” osmotic perturbation rather than very fast deformations as those simulated in this study. Hence, although such a mechanism should be taken into account when studying the evolution of axonal injury, it is here deemed not to affect our considerations.

Further method specific limitations have been previously addressed for both the axonal model and for molecular simulations of membranes.^26,113^ What we would like to address at this stage are possible alternatives to the methodology that was presented here to bridge the axon finite element and membrane molecular descriptions. An alternative to our approach could have been to describe also the lipid bilayer at the continuum (axonal) level and derive tensions (pressures) to be subsequently input into the molecular system. However, extracting continuum properties from molecular systems is a nontrivial problem and underlies several assumptions.^114–118^ Equating the strains at the two different length scales, on the contrary, allowed us to prescind from a continuum description of the lipid bilayer that would have been otherwise affected by several other assumptions. Another option is to directly apply strain rates (1, 10, and 40 1/s) to the molecular-based membrane model. However, at the moment, it is computationally unfeasible to obtain results at such time scale in a reasonable time frame. Applying local strain directly to the simulation allowed us to overcome the time scale issue.

The current models were used in cascade to investigate the response to a purely uniaxial stretch injury. Although this can be comparable to a controlled in vitro loading mode, it does not directly correlate with the in vivo scenario. Strains in the brain are in fact represented by a 3D tensor. Current finite element head models resolve, with slightly different methods, this tensor in the fiber direction and use this one-dimensional quantity as injury predictor.^119–122^ In future studies, the uniaxial condition could be dropped to study how an axonal model embedded in a matrix responds to a complete strain tensor. As a result of the different boundary conditions the axonal model and its axolemma might undergo deformations which are different from those observed in the present study.

## 6 Conclusions

In this study we successfully applied a top-to-bottom approach to better characterize the onset of axonal injury. By bridging continuum and molecular descriptions, we have provided quantitative evidences showing that mechanoporation of the axolemma is an event that cannot be excluded in a typical axonal injury scenario. In addition, we showed that pre-poration strain levels were characterized by an increased water permeability and interdigitated states. All in all, our study results provide increased knowledge of the axonal injury mechanism and have therefore potential positive implications for the development or optimization of neuroprotective measures targeting membrane integrity in the treatment of axonal injuries and brain injuries in general.

## 7 Acknowledgments

The authors thank the Swedish Research Council (VR-2016-05314), the European Union Horizon 2020 Research and Innovation Framework Programme under the Marie Skłodowska-Curie (grant agreement No. 642662), and the Swedish National Infrastructure for Computing (SNIC2017-1-491 and SNIC2018-3-548) for the support.

## 8 Author Contribution statement

AM, MS, AV and SK designed the study. AM performed and analyzed the finite element simulations, while MS and AV performed and analyzed the molecular dynamics simulation. AM, MS and AV prepared the manuscript. All authors critically reviewed the manuscript.

