## Supplementary material for "Localized axolemma deformations suggest mechanoporation as axonal injury trigger"

### Axon FE model

#### Microtubules:

**Morphological choices**

As now reported in the main text of the manuscript, given the axon cross-section and the proposed MTs densities (**Bray and Bunge 1981, Fadic et al. 1985, Malbouisson et al. 1985**) 19 rows of MTs were included in the axon RV, each containing two randomly placed discontinuities. The same approach was used in **Peter and Mofrad (2012), Lazarus et al. (2015), Soheilypour et al. (2015)**. Mechanical properties of the MTs have been very well identifies by previous studies.

**Material properties**

These filaments represent the stiffest part of the cytoskeleton and the Youngs' modulus (E=830 MPa) was derived by **Zhang et al. (2014)** with a numerical approach and was in extremely good agreement with experiments by **de Pablo et al (2003)** where MTs are locally probed. In the latter study, older bending experiments (which estimate a modulus E=1 to 1.2 GPa) were considered to overestimate the material properties of MTs because they usually probe length scales much longer than the one necessary to assess material properties. Being these filaments far stiffer then the remainder of our model, it can be said that a slightly stiffer modulus would not affect the localization of strains on the membrane.

#### Tau proteins:

**Morphological choices**

Cross-connection between microtubules (MTs) and between neurofilaments (NFs) and MTs were quantified in a seminal study by **Hirokawa (1982)**. There, MTs crosslinks were found to be less abundant than those between neurofilaments, which were reported having a 30-50 nm spacing. Inspecting the microscopy images in the same publication, a spacing of 120 nm was chosen for our model. Previous studies have also shown, through a sensitivity study, that tau protein density does not influence the energy associated with MTs stretching.

**Material properties**

Compared to the MTs, the material properties of these filaments are far less studied. The properties that we chose are based on previous studies by **Ahmadzadeh et al (2014,2015).** In these studies the viscoelastic characteristic of these cross-links was derived from previous experimental studies. While the spring stiffness is in the range of previously tested motor proteins (kinesin stiffness = uN/m ), we agree that its viscous component is the biggest assumption behind our model, which would however would affect mostly the microtubule deformation (still below failure ranges) rather than the onset of strain localization on the membrane. When working on the validation for the previous study however, we realized that increasing this value would have also led to unrealistic stiffnesses. At this stage unfortunately no additional experimental data is present regarding these proteins.

### Neurofilament Network:

#### Morphological choices

In the axon, neurofilaments run parallel the MTs array and their quantity has been reported to be 10 times that of MTs (**Wong et al. (1995)**). These intermediate filaments are considered to determine the axonal diameter since they form a very organized network, whose mesh-size is determined by the ~23 nm long side-arms, which present a 20 to 30-50 nm spacing (**Beck et al. (2009), Hirokawa (1984), Okonkwo et al. (1998)**). These quantities were used to create our discrete beam-meshed NFs network.

#### Material properties

Studies measuring the bulk properties of NF network in vitro revealed its viscoelastic behavior. However they could not replicate the extremely organized arrangement that this network has in vivo and their cross-linking system. Therefore the properties where obtaine dthrough calibration of the model against axon compression data by **Ouyang et al (2013)**.


**Bray, D. and Bunge, M.B., 1981. Serial analysis of microtubules in cultured rat sensory axons. *Journal of neurocytology*, *10*(4), pp.589-605.**

**Fadić, R., Vergara, J. and Alvarez, J., 1985. Microtubules and caliber of central and peripheral processes of sensory axons. *Journal of Comparative Neurology*, *236*(2), pp.258-264.**

**Malbouisson, A.M.B., Ghabriel, M.N. and Allt, G., 1985. Axonal microtubules: a computer-linked quantitative analysis. *Anatomy and embryology*, *171*(3), pp.339-344.**

**Peter, S.J. and Mofrad, M.R., 2012. Computational modeling of axonal microtubule bundles under tension. *Biophysical journal*, *102*(4), pp.749-757.**

**Lazarus, C., Soheilypour, M. and Mofrad, M.R., 2015. Torsional behavior of axonal microtubule bundles. *Biophysical journal*, *109*(2), pp.231-239.**

**Soheilypour, M., Peyro, M., Peter, S.J. and Mofrad, M.R., 2015. Buckling behavior of individual and bundled microtubules. *Biophysical journal*, *108*(7), pp.1718-1726.**

**Zhang, J. and Wang, C., 2014. Molecular structural mechanics model for the mechanical properties of microtubules. *Biomechanics and modeling in mechanobiology*, *13*(6), pp.1175-1184.**

**de Pablo, P.J., Schaap, I.A., MacKintosh, F.C. and Schmidt, C.F., 2003. Deformation and collapse of microtubules on the nanometer scale. *Physical review letters*, *91*(9), p.098101.**

**Hirokawa, N., 1982. Cross-linker system between neurofilaments, microtubules and membranous organelles in frog axons revealed by the quick-freeze, deep-etching method. *The Journal of cell biology*, *94*(1), pp.129-142.**

**Ahmadzadeh, H., Smith, D.H. and Shenoy, V.B., 2014. Viscoelasticity of tau proteins leads to strain rate-dependent breaking of microtubules during axonal stretch injury: predictions from a mathematical model. *Biophysical journal*, *106*(5), pp.1123-1133.**

**Ahmadzadeh, H., Smith, D.H. and Shenoy, V.B., 2015. Mechanical effects of dynamic binding between tau proteins on microtubules during axonal injury. *Biophysical journal*, *109*(11), pp.2328-2337.**

**Wong, P.C., Marszalek, J., Crawford, T.O., Xu, Z., Hsieh, S.T., Griffin, J.W. and Cleveland, D.W., 1995. Increasing neurofilament subunit NF-M expression reduces axonal NF-H, inhibits radial growth, and results in neurofilamentous accumulation in motor neurons. *The Journal of cell biology*, *130*(6), pp.1413-1422.**

**Beck, R., Deek, J., Jones, J.B. and Safinya, C.R., 2010. Gel-expanded to gel-condensed transition in neurofilament networks revealed by direct force measurements. *Nature materials*, *9*(1), p.40.**

**Hirokawa, N., Glicksman, M.A. and Willard, M.B., 1984. Organization of mammalian neurofilament polypeptides within the neuronal cytoskeleton. *The Journal of cell biology*, *98*(4), pp.1523-1536.**

**Hui, O., Nauman, E. and Shi, R., 2010, June. Contribution of cytoskeletal elements to the mechanical property of axons. In *2010 18th Biennial University/Government/Industry Micro/Nano Symposium* (pp. 1-5). IEEE.**

### Sensitivity study

A sensitivity study was performed with one of the ten geometries to assess which material properties affect cortex deformation the most. Keeping in mind that the original properties yielded an axonal stiffness comparable to experimental data, the following parameters sets were selectively changed by increasing or reducing them by 100%:

- **Tau viscosity (mu)** - in this case the degenerate case of purely elastic tau proteins was also included
- **Cortex properties (G0,G_inf and Beta)**
- **Neurofilament (K, mu)**

The results (**Figure S1**) show that both the cortex and neurofilaments properties minimally afftect the deformation in the cortex in the range of interest. Increasing/reducing tau viscosity has almost no effect at strain rate 1 /s, whereas an effect is visible at strain rate 10/s mostly before the average poration threshold, which is indicated with a black line. At strain rate 40 /s the effect of a change in viscosity is more pronounced. Doubling or halving the viscosity respectively “delays” or “anticipates” of 1% axonal strain the crossing of the mechanoporation limit: The curved are reunited at 18% axonal strain (not shown).

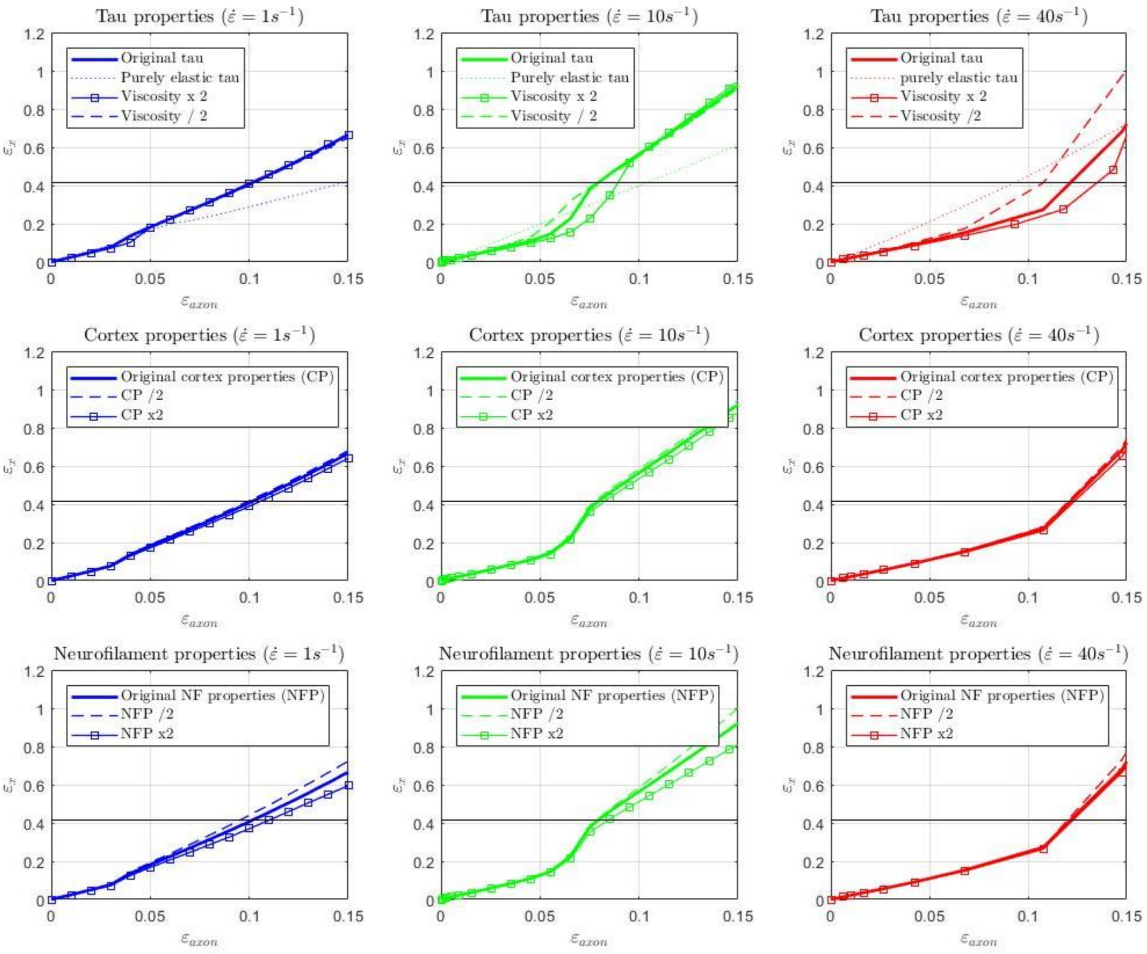


**Figure S1** Maximum 1st Principal Green-Lagrange strain in the cortex as a result of changes in tau protein viscosity (first row), cortex properties (CP, second row), neurofilaments properties (NP, second row). First, second, and third column show the results for strain rate 1, 10, and 40 /s.

| Membranes Weight Fraction | *Phosphatidyl choline* | *Phosphatidyl ethanolamin e* | Sphingomyelin | Phosphatidylserine | *Cholesterol* | *Glycolipid* | *Other* |
|---|---|---|---|---|---|---|---|
| **Plasma membrane model** [**this study**] | 30 | 17 | 17 | 6 | 17 | 5 | 8 |
| **Liver cell plasma membrane** [**Alberts, B., 2017**] | 24 | 7 | 19 | 4 | 17 | 7 | 22 |
| **Red blood cell plasma membrane** [**Alberts, B., 2017**] | 17 | 18 | 18 | 7 | 23 | 3 | 14 |
| **Axolemma** [values range from **DeVries et al. (1981) (1999)** and **Zambrano et al. (1971)**] | 11-29 | 11-26 | 3-13 | 3-10 | 22-27 | 15-25 | 4-17 |

**Table S1** Comparison between the lipid composition (weight %) of the model used in this study against the available experimental data. Note that the lipid composition in manuscript is reported in mol %. Here we  report the lipid composition in weight % to compare with experimental data.


**Alberts, B. Molecular biology of the cell. (Garland science, 2017).**

**DeVries, G.H., Zetusky, W.J., Zmachinski, C. and Calabrese, V.P., 1981. Lipid composition of axolemma-enriched fractions from human brains. *Journal of lipid research*, *22*(2), pp.208-216.**

**DeVries, G.H., Campbell, B. and Saunders, R., 1999. Isolation and characterization of unmyelinated axolemma from bovine splenic nerve. *Journal of neuroscience research*, *57*(5), pp.670-679.**

**Zambrano, F., Cellino, M. and Canessa-Fischer, M., 1971. The molecular organization of nerve membranes. *Journal of Membrane Biology*, *6*(4), pp.289-303.**

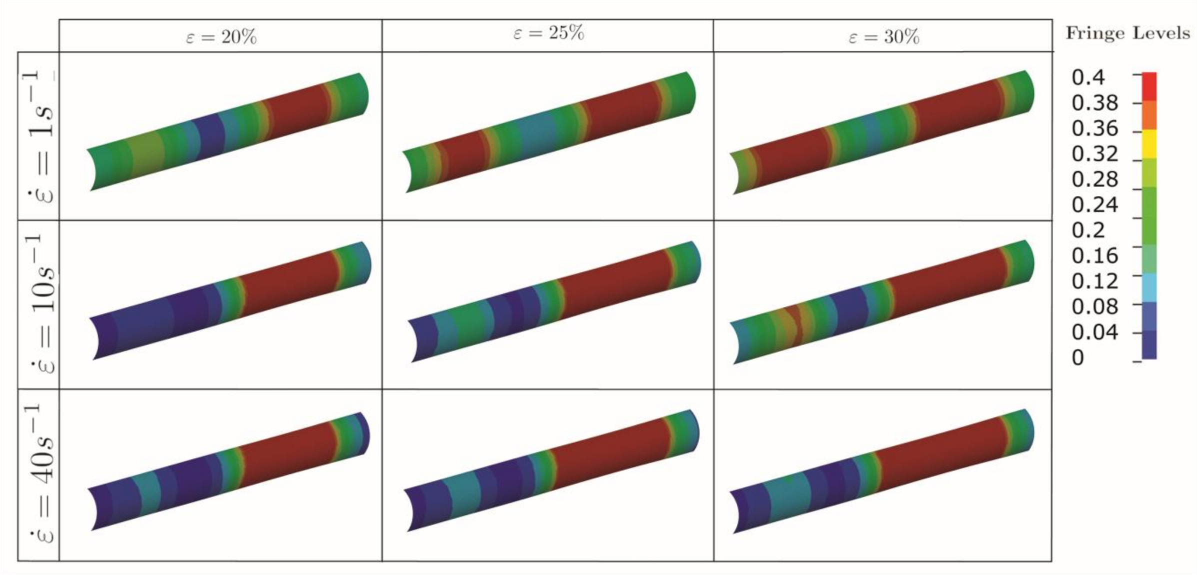


**Figure S2** Fringe plots showing the 1st Principal Green-Lagrange strain along the axonal cortex as a result of 20, 25, and 30 % axonal strains in the first, second, and third column, respectively. In the first, second, and third row results for strain rates of 1, 10, and 40 /s are reported.

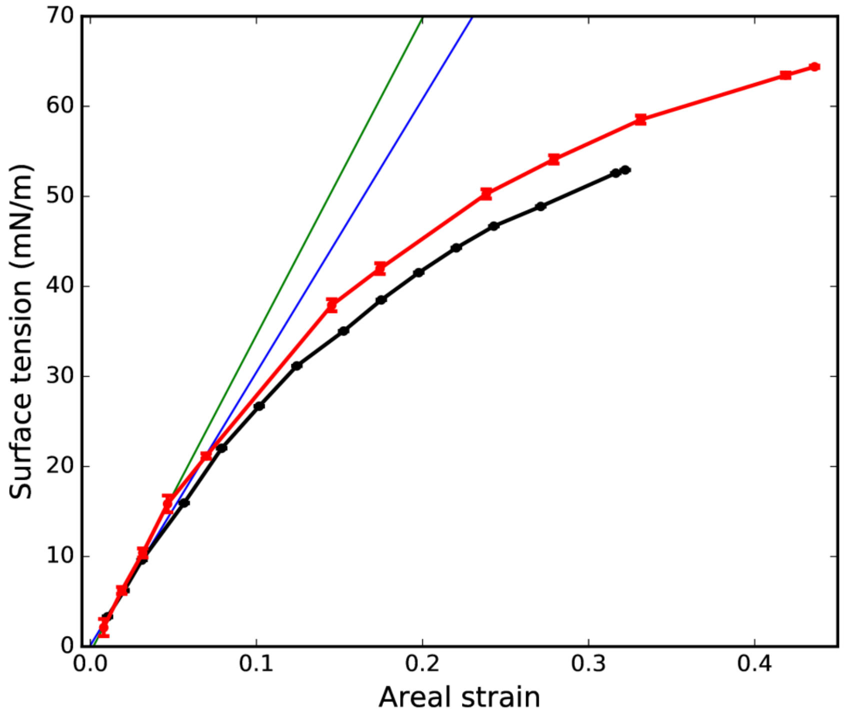


**Figure S3** Bilayer surface tension (γ) versus areal strain ($\varepsilon_A$). The equilibrium values for surface tension and bilayer surface area are used as reference. Area values have been calculated using Voronoi algorithm (Lukat et al. doi: 10.1021/ci400172g. ). Results for bilayers in absence and in presence of $Na_v1.1$ protein are reported in black and in red, respectively. The surface tension was averaged on the last 1 µs and reported errors are based on block average over 5 blocks. The slope of the curve corresponds to the bilayer area compressibility modulus, $K_A = \left(\frac{d\gamma}{d\varepsilon_A}\right)$. A value of $K_A$ of 303 mN/m (blue line, membrane system) and of 352 mN/m (green line, membrane-protein system), was obtained by linear regression of the first three points (at $\varepsilon_A < 0.05$).

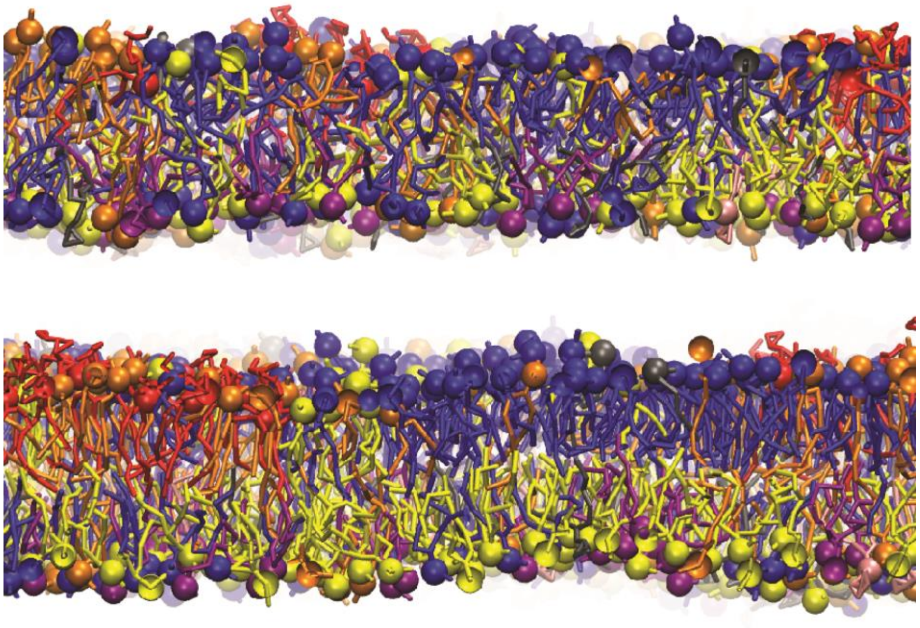

**Figure S4**

Lipid interdigitation. Top panel: snapshot at 550 ns for lipid bilayer at $\varepsilon_x$ =0.32 (lipid tails are interdigitated). Low panel: snapshot at 5000 ns for lipid bilayer at equilibrium (no lipid tail interdigitation). PC lipids are in blue, PE in yellow, SM in orange, PS in purple, GM in red, anionic lipids in pink, and the rest of lipids in gray, water and cholesterol molecules are not visualized for clarity. Lipid head group beads are shown in ball presentation for clarity.

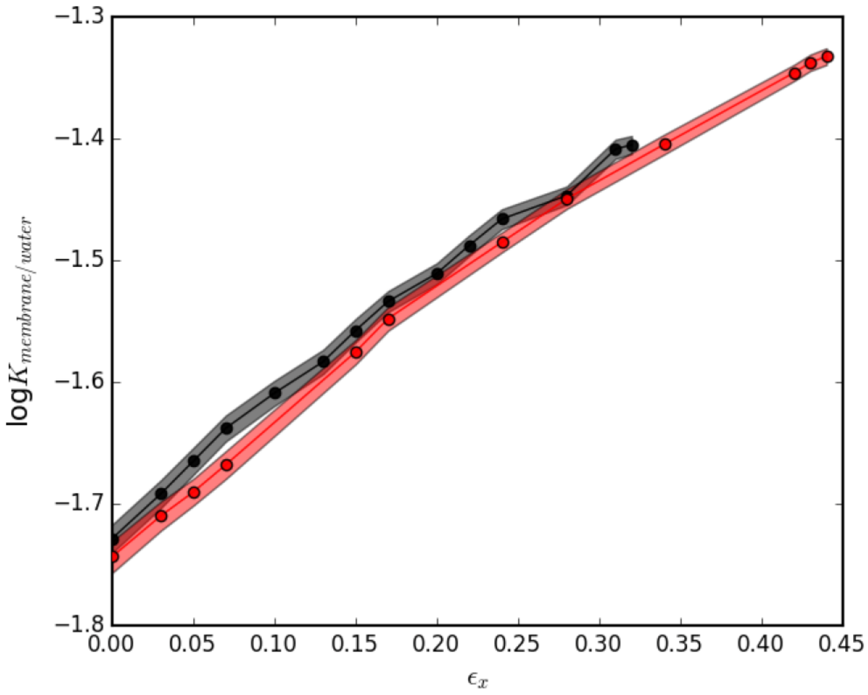


**Figure S5** Water partition coefficient between aqueous solution and lipid bilayer as a function of bilayers local strain in absence (black) and in presence of $Na_v$ 1.1 protein ( red). The shaded area represents the standard deviation.

**Movie S1**: Bilayer rupture at strain $\varepsilon_x$ =0.34 (160 ns)

https://drive.google.com/open?id=1ho2sqKSk7OcUhlxAsA801D1Fuec-k44U

**Movie S2**: Bilayer rupture in presence of $Na_v$1.1 at strain $\varepsilon_x$ =0.47 (40 ns)

https://drive.google.com/open?id=1CTK_xtw_OW-IZlcjME6JapWywOg7GJlG
